# Evaluation of Kernel Low-Rank Compressed Sensing in preclinical Diffusion Magnetic Resonance Imaging

**DOI:** 10.1101/2023.02.09.527467

**Authors:** Diego Alves Rodrigues De Souza, Hervé Mathieu, Jean-Christophe Deloulme, Emmanuel L. Barbier

**Author notes:** Co-Last authors (Jean-Christophe Deloulme, Emmanuel L. Barbier). Corresponding author, Emmanuel L. BARBIER, Grenoble Institut des Neurosciences – U1216, Team «Functional Neuroimaging and Brain Perfusion», Chemin Fortuné Ferrini, 38700 La Tronche.

## Abstract

Compressed Sensing (CS) is widely used to accelerate clinical diffusion MRI acquisitions, but it remains under-utilized in preclinical settings. In this study, we optimized and compared several CS reconstruction methods for diffusion imaging.

Different undersampling patterns and two reconstruction approaches were evaluated: conventional CS, based on Berkeley Advanced Reconstruction Toolbox (BART-CS) toolbox, and a new Kernel Low-Rank (KLR)-CS, based on Kernel Principal Component Analysis and low-resolution-phase maps. 3D CS acquisitions were performed at 9.4T using a 4-element cryocoil on mice (wild type and a *MAP6* knockout). Comparison metrics were error and Structural Similarity Index Measure (SSIM) on fractional anisotropy (FA) and mean diffusivity (MD), as well as reconstructions of the anterior commissure and fornix. Acceleration factors (AF) up to 6 were considered.

In the case of retrospective undersampling, the proposed KLR-CS outperformed BART-CS up to AF=6 for FA and MD maps and tractography. For instance, for AF=4, the maximum errors were respectively 8.0% for BART-CS and 4.9% for KLR-CS, considering both FA and MD in the corpus callosum. Regarding undersampled acquisitions, these maximum errors became respectively 10.5% for BART-CS and 7.0% for KLR-CS. This difference between simulations and acquisitions arose mainly from repetition noise, but also from differences in resonance frequency drift, signal-to-noise ratio, and in reconstruction noise. Despite this increased error, fully sampled and AF=2 yielded comparable results for FA, MD and tractography, and AF=4 showed minor faults. Altogether, KLR-CS based on low-resolution-phase maps seems a robust approach to accelerate preclinical diffusion MRI.

## 1. Introduction

Diffusion MRI, based on diffusion-weighted imaging (DWI), is used in neuroscience research to characterize anatomical connectivity^1^. At the preclinical level, anatomical connectivity mapping helps to characterize transgenic animal models used in various areas, from neurodevelopment^2^ to neurodegeneration^3,4^. To obtain reliable connectivity data, it is now accepted that at least 30 diffusion directions are required; the precise number depends on the analysis approach^5^. Advanced diffusion models require even richer datasets (more diffusion directions and/or use of several diffusion gradient values) and therefore take longer to acquire^6,7^. Currently, these in vivo acquisitions can be used for humans owing to several well-known acquisition strategies: parallel imaging^8,9^, compressed sensing (CS)^10^, and multiband acquisition^11,12^.

In preclinical settings, however, these developments are not routinely available, despite several proof-of-principle studies. Moreover, the small voxel sizes required to map tracts in mouse or rat brains (100 μm or less) contribute to further increasing the DWI acquisition times, which are much longer than in clinical settings^7,13^. Therefore, most preclinical studies are performed ex vivo with acquisition durations that routinely exceed 8h^6,14–16^. Such long acquisition times may decrease image quality because of hardware and/or sample drift^17^ and can limit the number of samples used owing to cost and/or scan time availability.

One of the most attractive means to reduce acquisition duration is CS^10,18–20^, which has recently been evaluated in mouse brain connectomics^6,21^. The Johnson group reported connectivity data obtained with an acquisition pattern based on Monte-Carlo sampling, which was constant across diffusion directions^6,16^. Acquired data were reconstructed using the SparseMRI CS approach developed by Lustig et al^10^, and the impact of reducing the scan time on mean diffusivity (MD), fractional anisotropy (FA), and on the ability to detect white-matter tracts and fiber-crossings was evaluated^16^. Their approach was also evaluated in a model of autism spectrum disorders^6^. In their settings, an 8-fold reduction in scan time (from 92.6h to 11.8h) was acceptable, but the scan time remains long. More recently, Zhang et al proposed a kernel low-rank (KLR) method to jointly reconstruct multiple diffusion-weighted images^21,22^, using undersampling patterns that vary with the diffusion direction^20^. In their settings, the KLR-CS appears more efficient than the conventional CS proposed by Lustig et al for a 5.56-fold reduction in scan time (from 8h to 1.4h)^10^. However, the proposed KLR approach has been evaluated on magnitude-filtered data, i.e. data in which the Hermitian symmetry in k-space has been enhanced, and using retrospective undersampling^21^. Altogether, to our knowledge, there is no implementation of a KLR-CS approach able to efficiently handle complex undersampled DWI acquisitions.

In this study, we thus evaluated both the acquisition, using a four-channel receive coil, and a KLR-CS reconstruction protocol adapted to handle complex undersampled DWI data. Data were acquired on a preclinical mouse model that exhibits altered brain connectivity: the *MAP6* knockout model^23^. The DWI undersampling parameters were optimized using simulations. A KLR-CS approach, based on the use of low-resolution-phase (LRP) maps, was introduced, and parametric maps and fiber tracts from undersampled acquisitions were analyzed, using the Berkeley Advanced Reconstruction Toolbox (BART)-CS as reference. Finally, differences between simulations and acquisitions were explored, considering the change in signal-to-noise ratio, frequency drift, and repetition noise.

## 2. Material and Methods

### 2.1 Animals

The study protocol was approved by the local animal welfare committee (Comité Local GIN, C2EA-04 - APAFIS number 21234-2019031308592774) and complied with EU guidelines (Directive 2010/63/EU). Every precaution was taken to minimize the number of animals used and the stress on animals during experiments. The *MAP6*-deficient mouse line used in this study is on the C57BL/6 genetic background^24^. Adult mice that were heterozygous for *MAP6* (*MAP6*^+/-^) and their wild-type (WT) littermates were obtained by crossing *MAP6^+/-^* mice.

As the complete *MAP6* knockout (*MAP6*^+/-^) leads to major changes in brain connectivity^23^, we used the partial (heterozygous *MAP6*^+/-^) knockout to achieve subtle changes in tract morphology. Thus, we challenged the level of detail quantifiable by CS reconstructions.

### 2.2 Brain preparation for ex vivo MRI acquisitions

Brains were prepared according to a previously reported protocol^15^. After transcardiac perfusion with a 4% paraformaldehyde solution containing Gd-chelate (6.25 mM of Gd-DOTA; Guerbet Laboratories, Roissy, France), mice were decapitated. After the removal of skin and muscles, the skull was immersed in the same fixing solution for 4 days and transferred into Fomblin oil (FenS chemicals, Goes, Netherlands), an oil that generates no MRI signal but has a magnetic susceptibility close to that of brain tissue. MRI was performed at least 7 days after brain fixation. This protocol ensures a homogeneous distribution of the Gd-DOTA throughout the whole mouse brain^25^. The contrast agent decreased brain T1 from 1000±102 to 110±13 ms and brain T2 from 27.3±3.1 to 18.3±2.6 ms (data not shown), allowing a reduction in acquisition times.

### 2.3 Optimization of the CS undersampling pattern for DWI

To select an undersampling approach, 20 different k-space undersampling patterns were explored, by retrospectively undersampling the FS DWI datasets acquired from the 3 WT brains and using the methods described below:

- Monte-Carlo vs Poisson-disc sampling. Monte-Carlo sampling patterns were generated through a variable probability density function with a flat region in its center^16,21^. Poisson-disc sampling patterns were generated using the algorithm embedded in BART. For this pattern, we evaluated regular and elliptical sampling as well as uniform and variable sampling density. To ensure fair comparisons, the area of the FS center of k-space was kept constant across sampling patterns;
- Single-vs multi-mask sampling^26^. The single-mask approach stands for the use of the same undersampling pattern for all DWI volumes and the multi-mask approach for the use of different undersampling patterns for each DWI volume;
- FS vs undersampled b0 images. To ensure a comparable global AF across the whole DWI datasets (b0 + diffusion direction images), the AF per diffusion direction (AFdiff) was increased in the case of FS b0 images. The behavior of AF and AFdiff, as a function of the sampling strategies for b0 images, is further described in Supp. Fig. 1.

AF values of 2, 3, 4, and 6 were explored. Images were reconstructed and MD and FA were computed across the whole brain. The most robust undersampling strategy was determined based on the median error and SSIM (see “2.7 Comparison metrics”).

### 2.4 Image acquisitions

MRI acquisitions were carried out at 9.4 T (Biospec Avance III HD, Bruker, Ettlingen, Germany; IRMaGe facility) using a 4-channel head surface cryocoil for reception, a volume coil for transmission, and Paravision 7. The following sequences were performed:

- a 3D CS T1w gradient-echo MRI acquisition: repetition time (TR)=50.0 ms, echo time (TE)=6.61 ms, flip angle=40°, signal accumulations=4, field of view (FOV): 18 × 12.8 × 12.8 mm^3^, isotropic spatial resolution: 100 μm^3^ (180×128×128 matrix size), acceleration factor (AF)=2. The acquisition duration was 23 min. These anatomical data were used to overlay an atlas for the DWI analysis (see section 2.6 ‘Tractography’).
- a 3D CS, spin-echo, diffusion-weighted images (DWI): TR=150.0 ms, TE=19.0 ms, flip angle=90°, signal accumulations=1, FOV: 18×9×11 mm^3^, isotropic spatial resolution: 100 μm^3^, δ=4.0 ms, Δ=9.6 ms, bandwidth=45454.5 Hz. 33 separate volumes were acquired: 3 baseline images (b0 images) acquired without diffusion weighting (b~0 s/mm^2^) and 30 diffusion directions (b=3000 s/mm2). The gradient directions were generated using an optimal distribution across the surface of the sphere^27^. The acquisition duration of the fully sampled (FS) dataset was 13h37min. This sequence allows changing the 3D undersampling pattern for each diffusion direction (code available for download at https://github.com/nifm-gin/compressedSensing). Several acquisitions were performed with different AF, as summarized in Supp. Table 1. To estimate and limit the impact of the frequency drift, the main resonance frequency was calibrated before and after each DWI acquisition.

### 2.5 Image reconstruction

Data were reconstructed using the Matlab environment (MATLAB2021b; The MathWorks, MA, USA). For the undersampled dataset, two reconstruction approaches were performed:

- Conventional CS reconstructions were performed using the BART^28,29^ toolbox v0.7.00 (https://mrirecon.github.io/bart/), which calculates coil sensitivity maps using ESPIRiT calibration^30^. For the minimization process, we used the parameters optimized in^16^: λ_1_ = 0.005, λ_2_ = 0.002, max iteration =200. This toolbox was chosen to have the same reference technique as in the original KLR-CS paper^21^.
- KLR-CS reconstructions of diffusion-weighted images were performed using an in-house adaptation of the original KLR toolbox as published^21,22^. However, the original reconstruction method was evaluated on data after magnitude filtering, a step that enhances the Hermitian symmetry of k-space^21^. Therefore, this method – hereafter called “magnitude-filtered KLR-CS” – is not designed for acquired undersampled datasets as the data-consistency step is no longer efficient. We therefore proposed and evaluated two adaptations of the original KLR toolbox: a composite kernel principal component analysis (KPCA) adapted from the composite real PCA of complex signals^31^ and LRP maps. For the KLR-CS based on composite KPCA, we replaced the original input to the KPCA module — i.e. magnitude data — with composite data, which is a concatenation of the real and imaginary parts of the data. The output of composite KPCA is then reorganized to produce complex data before enforcing data consistency. For the KLR-CS based on LRP maps, the phase information related to the FS center of k-space (i.e. the low-resolution part of the dataset) is stored before the self-training step and added back to the output magnitude image, yielding a complex image that can then be used to enforce data consistency. This approach using the phase information of the entire undersampled k-space was also tested but led to lower quality results. For KLR training, the low-resolution, 4-channels, 4D images obtained from the FS region at the center of the k-space were used.

In addition, all the FS DWI datasets (n=6 brains; 3 WT and 3 *MAP6*^+/-^) were retrospectively undersampled, using the same undersampling pattern and AF values as the one used for the acquisitions, and reconstructed with the two methods described above. Thereby, metrics obtained from simulated (i.e. retrospective) undersampling could be compared to that obtained from undersampled acquisitions.

### 2.6 MD and FA maps and tractography

DWI data was processed using MRtrix (v3.0_RC3-137, https://www.mrtrix.org/). After image denoising, unringing, and brain masking, tensor-derived FA and MD maps were generated. FA and MD were measured in the whole brain and two regions of interest (ROI), corpus callosum and caudate putamen.

The fiber orientation distributions were computed with constrained spherical deconvolution (CSD)^32,33^ and default parameter values (for instance, maximum harmonic order lmax = 4^34^). Whole brain tractography was computed with a probabilistic streamline tracking^35,36^ (iFOD2 algorithm) and three seeds per voxel. Empirically optimized tracking parameters on our mouse brain data were applied after normalization of the fiber orientation distributions: step size = 0.01 mm, radius of curvature = 0.07 mm ^37^, min length = 0.1 mm, max length = 50 mm ^38^ and cutoff=0.2. In addition, several major tracts were isolated as follows. For each tract, several ROIs were defined, according to a mouse brain atlas^39^. A ROI was a 2D surface in one orientation (coronal, horizontal, sagittal). Each ROI-tract association then received either the label ‘AND’ or ‘NOT’. The fibers of one particular tract were selected as all the fibers that cross the ROIs labeled ‘AND’ minus the fibers that cross the ROIs labeled ‘NOT’. Supp. Table 2 shows the list of ROIs, with their labels and the orientation in which they were considered, for each tract evaluated in this study.

### 2.7 Comparison metrics

To compare parametric maps, two metrics were used. The absolute error (%) — called “error” in the rest of the manuscript — was computed voxel-wise using:

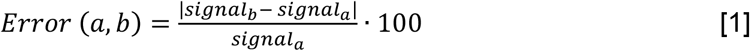

where signala and signalb are the values of a given voxel in the reference image and the reconstructed image, respectively. The Structural Similarity Index Measure (SSIM, ranging from 0 (worst) up to 1 (best)) was computed as^40^:

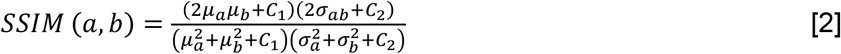

where μ_a_, μ_b_, σ_a_, σ_b_, and σ_ab_ are the local means, standard deviations, and cross-covariance for the reference and the reconstructed images, respectively. C_1_ and C_2_ were set to 10^-4^ and 9.10^-4^ (default Matlab values).

Both metrics were calculated for each reconstruction, considering the whole 3D brain. Error and SSIM maps were obtained using the FS dataset as the reference, unless mentioned otherwise. In addition, to allow tractogram comparisons, the mean fiber length was computed. To remove the image shift caused by the frequency drift that occurs during hours-long acquisitions, raw DWI images were registered to the FS images using an in-house rigid registration tool prior to computing error and SSIM metrics. Across each brain, the median error and the median SSIM were retained, to limit the effect of locally high errors such as the ones observed on the edges or in moving structures (ventricles, vessels).

Statistical analyses were performed in GraphPad Prism version 9.4.1 for Windows (GraphPad Software, La Jolla California USA, www.graphpad.com). The *p*-values mentioned along this article correspond to paired *t*-tests, unless stated otherwise.

## 3. Results

### 3.1 Optimization of the CS for DWI using simulations

Regarding the choice of undersampling patterns, examples based on multi-mask and Monte-Carlo or Poisson-disc sampling are shown in Supp. Fig. 2A, for different AF values. Overall, while the choices of Monte-Carlo multi-mask sampling clearly yielded better results (Supp. Fig. 2B-E), the full sampling of b0 images did not provide a major advantage. To ease comparison with the previous study^21^, we choose to keep that full sampling of b0 images. Altogether, the undersampling pattern based on Monte-Carlo with FS b0 images and multi-mask was used for all subsequent evaluations.

Using this sampling pattern, we then compared the effects of composite KPCA and LRP using complex data and AF between 2 and 6. For sake of comparison, we evaluated the original KLR-CS method using k-spaces from complex images instead of k-spaces from magnitude-only images. Supp. Fig. 3 shows FA and MD maps obtained from the two proposed KLR-CS, BART-CS, and the magnitude-filtered KLR-CS reconstructions. The corresponding error and SSIM maps are shown in Supp. Fig. 4A-B, E-F, respectively. As expected, BART-CS, LRP KLR-CS, and KPCA KLR-CS, which all handle complex data, yielded better results than the Magnitude-filtered KLR-CS. For example, for AF=2, the median error on FA with LRP KLR-CS was 4.79±0.12% while it was 9.69±0.12% with Magnitude-filtered KLR-CS (Supp. Fig. 4C-D, G-H). This confirms that the original, magnitude-filtered, KLR CS approach is not adapted to complex data. In the rest of the manuscript, we thus focus on LRP and KPCA KLR-CS. Regarding FA (Supp. Fig. 4C-D), the LRP KLR-CS outperformed BART-CS for AF=2 and AF=3 (*p*<0.0001) and KPCA KLR-CS for all AFs (*p*<0.0001). For AF=4 and 6, LRP KLR-CS had similar results to BART-CS. Regarding MD (Supp. Fig. 4G-H), LRP KLR-CS outperformed BART-CS and KPCA KLR-CS for all AFs (*p*<0.01). Therefore, LRP KLR-CS, hereafter called “KLR-CS” for simplicity, was used to reconstruct all the data reported in the subsequent studies.

Fig. 1A,B shows examples of FA and MD maps obtained from a WT mouse as a function of AF using BART-CS and KLR-CS reconstructions, optimized as described. There were no obvious faults observed in the brain slices. The focus on the hippocampus shows decreased contrast as AF increases, in particular for BART-CS reconstructions. For quantitative analysis, the corresponding error and SSIM maps are shown in Fig. 2A,D, and the median error and median SSIM values across the six brains are shown in Fig. 2B,C for FA and Fig. 2E,F for MD. Since the median error and median SSIM of FA and MD maps were not significantly different between the WT and *MAP6*^+/-^ groups (unpaired *t*-test, *p*>0.05), data were pooled to improve statistical power in the methodological study. As AF increased from 2 to 6, there was an increase in median error together with a decrease in median SSIM. Overall, KLR-CS reconstructions led to smaller median errors and higher median SSIM than BART-CS, regardless of AF. For instance, with AF=2 and for FA maps, BART-CS and KLR-CS reconstructions had a median error of 5.78±0.11 and 4.79±0.12% (*p*<0.0001) and a median SSIM of 0.976±0.001 and 0.982±0.001 (*p*<0.0001), respectively. For MD maps under AF=2, BART-CS and KLR-CS had a median error of 1.41±0.04 and 0.71±0.02% (p<0.0001) and a median SSIM of 0.988±0.001 and 0.997±0.001 (*p*<0.0001), respectively. Altogether, the KLR-CS reconstructions were of very good quality up to AF=4, with an error below 2% and a SSIM above 0.970.

**Fig. 1:**
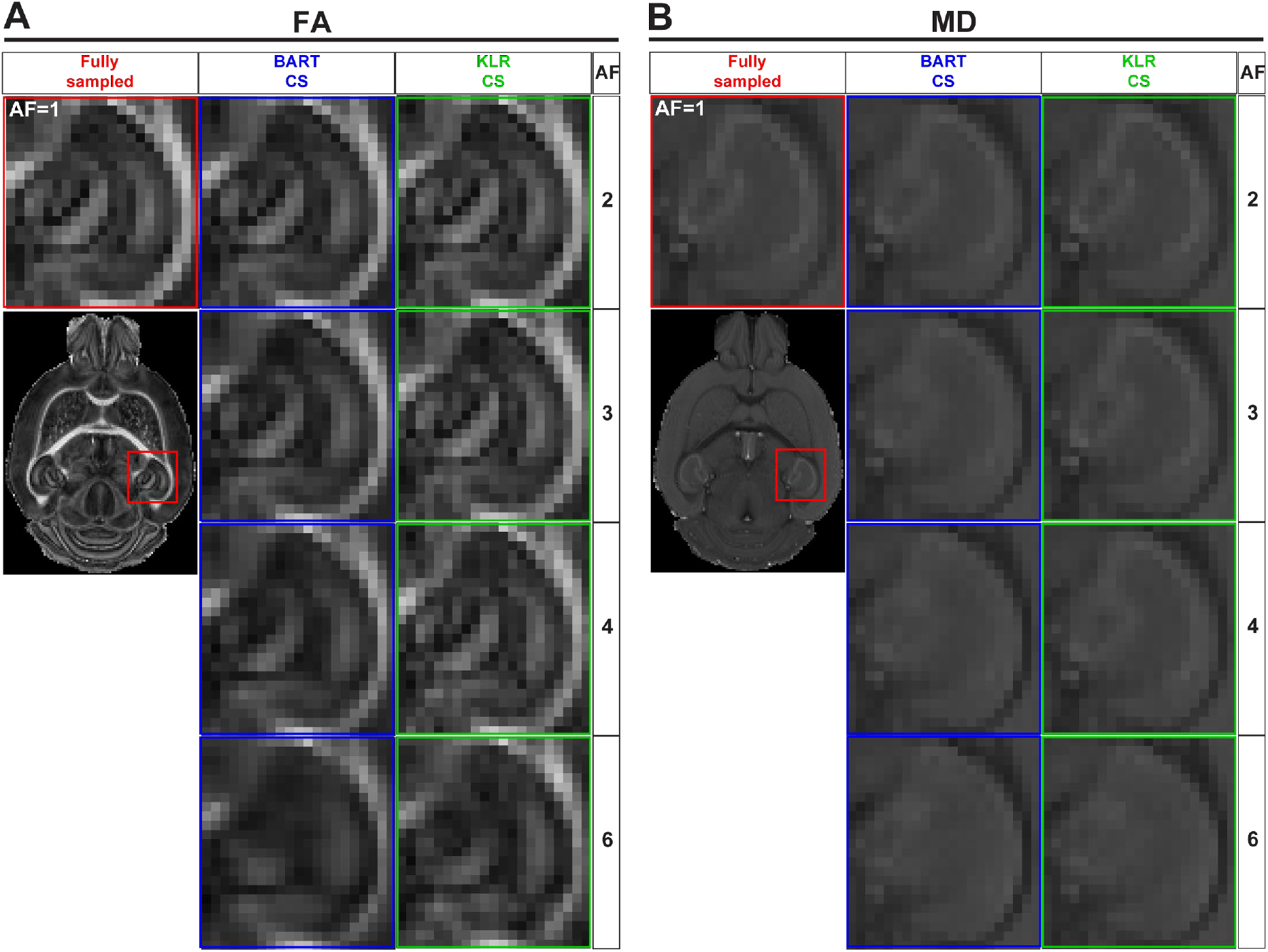
FA and MD maps from fully sampled data and after simulated undersampling, using BART-CS and KLR-CS reconstructions. Examples of (**A**) FA and (**B**) MD maps from a magnification on the hippocampus, obtained from fully sampled (red; AF=1), BART-CS (blue), and KLR-CS (green) reconstructions, using different AFs.

**Fig. 2:**
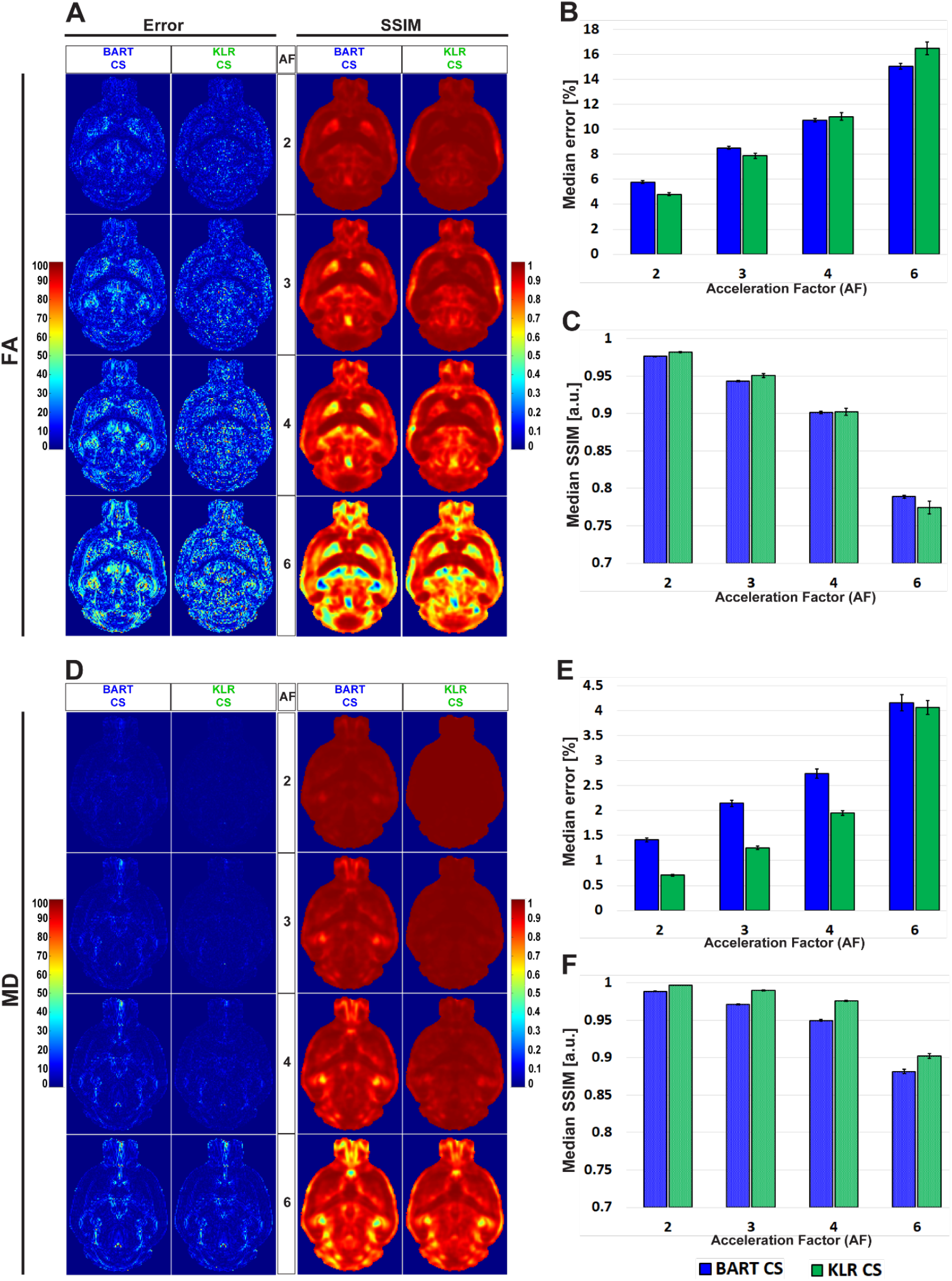
Error and SSIM, derived from FA and MD maps obtained after simulated undersampling and reconstruction by BART-CS and KLR-CS. Error and SSIM maps for (**A**) FA and (**D**) MD, using BART-CS and KLR-CS reconstructions, for different AF values. Corresponding (**B,E**) median error and (**C,F**) median SSIM obtained for the full 3D brain, using fully sampled data as the reference. Results are expressed as mean ± standard deviation across the animals (n=6).

Tractographies of the anterior commissure of WT and *MAP6*^+/-^ mice — a model for which we expect tract differences — are shown in Fig. 3 for different AF values. First, as expected, the WT and *MAP6*^+/-^ mice exhibited different mean fiber lengths, as observed on the FS data (unpaired *t*-test, *p*=0.0281) (Fig. 3C). For AF=2, 3 and 4 and with the parameters used for the tractography, the two reconstruction methods had similar performances (*p*>0.05 for AF=2, 3 and 4 vs AF=1) and maintained the ability to observe the reduction in mean fiber length between WT and *MAP6*^+/-^ mice. For AF=6, KLR-CS remained robust while BART-CS sometimes missed the anterior part of the anterior commissure in *MAP6*^+/-^ mice (Fig. 3B).

**Fig. 3:**
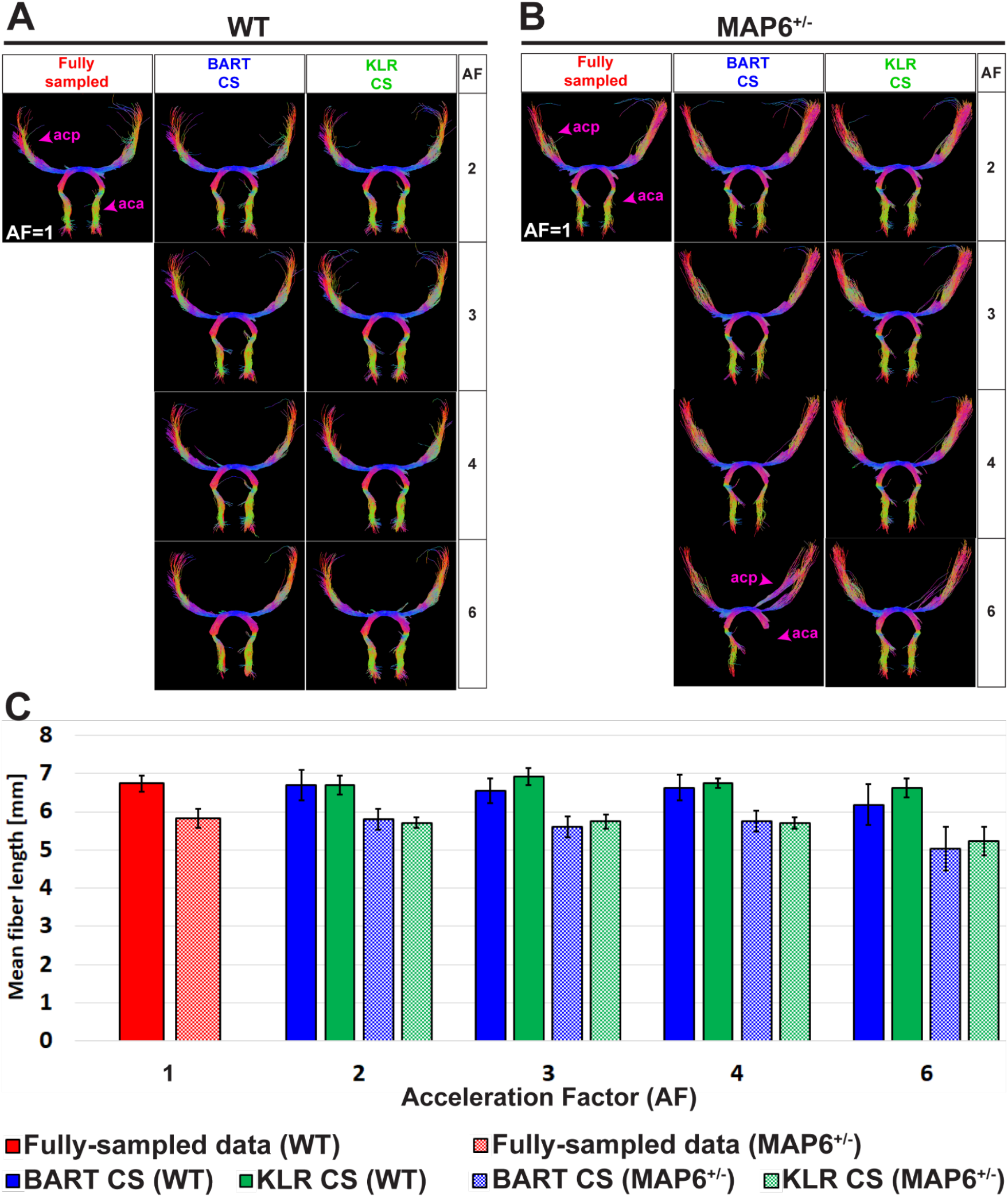
Tractography of the anterior commissure from fully sampled data and after simulated undersampling, using BART-CS and KLR-CS reconstructions. Anterior commissure (ac) of a (**A**) WT and a (**B**) *MAP6*^+/-^ mouse, derived from fully sampled (red) and, after simulated undersampling, using BART-CS (blue), and KLR-CS (green) reconstructions, for different AFs. (**C**) The corresponding mean fiber length of the ac. Results are expressed as mean ± standard deviation across 3 animals (3 WT or 3 *MAP6*^+/-^). acp: ac posterior part; aca: ac anterior part.

Overall, KLR-CS and BART-CS yielded comparable results for AF=2, 3, and 4, but KLR-CS remained reliable at AF=6 while BART-CS did not.

### 3.2 Evaluation of CS DWI acquisitions

After evaluating BART-CS and KLR-CS methods using simulated undersamplings, we reconstructed undersampled acquisitions carried out with our in-house DWI Spin-Echo CS sequence. AF=2 and 4 were used as they corresponded to the most promising simulation results. For the acquisitions, we evaluated the FA and MD maps and the ability to reconstruct tracts. No difference may be seen on the whole maps (not shown), however, magnifying the complex region of the hippocampus revealed reduced image contrasts and some spatial smoothing as AF increased (Fig. 4). Visually, this smoothing was less pronounced for KLR-CS than for BART-CS, in line with the simulation results. Corresponding error and SSIM maps (Fig. 5A-D) and values (Fig. 5E-H) confirmed the better performance of KLR-CS over BART-CS (e.g., for AF=2, MD error: 2.59±0.12% and 2.13±0.12%, *p*<0.0001; MD SSIM: 0.964±0.002% and 0.977±0.002, *p*<0.0001).

**Fig. 4:**
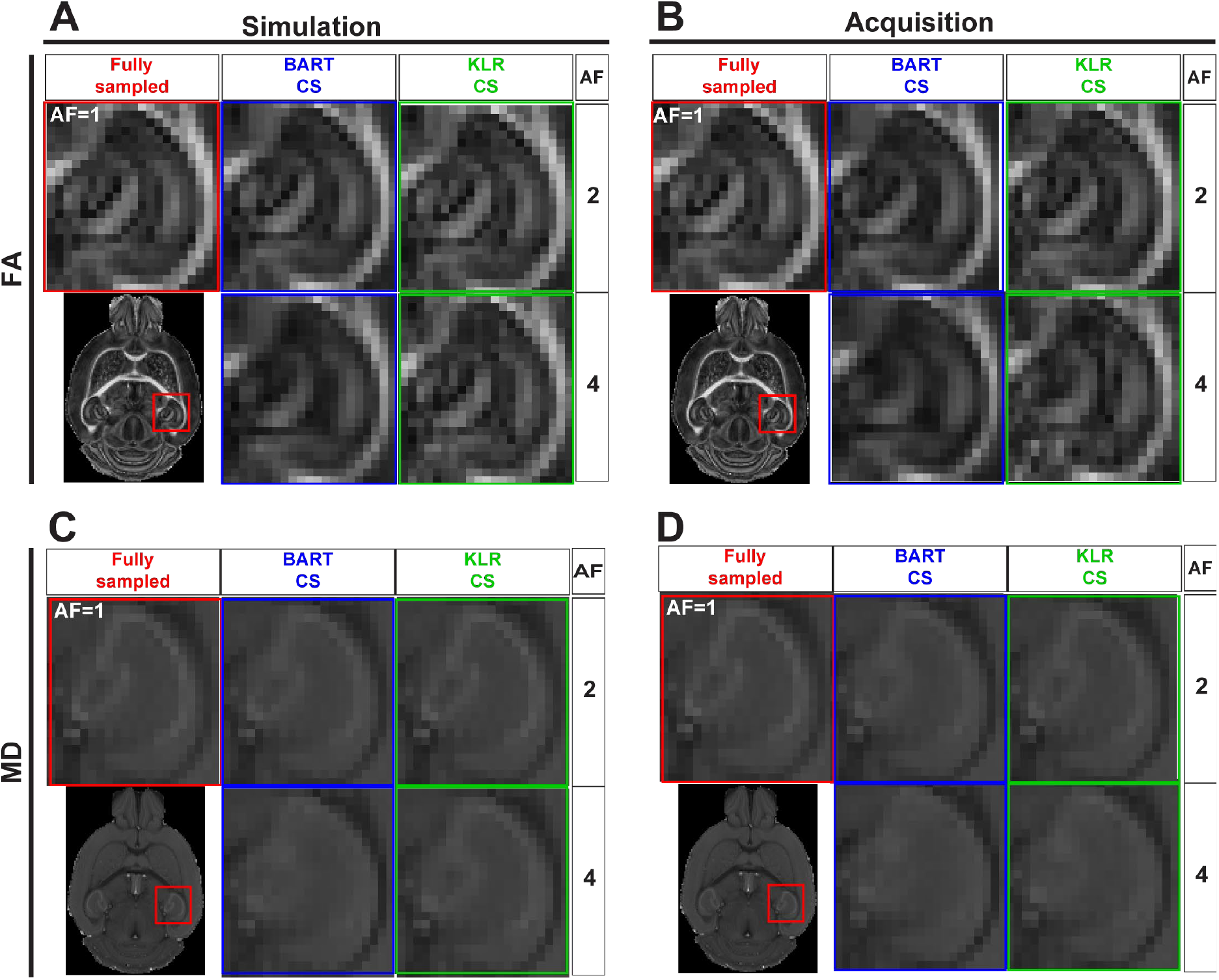
FA and MD maps from fully sampled data, after simulated undersampling, and for CS acquisitions, using BART-CS and KLR-CS reconstructions. (**A,B**) FA and (**C,D**) MD maps from a magnification on the hippocampus. Maps in the left column (**A,C**) are obtained after retrospective undersampling (Simulation) and maps in the right column (**B,D**) after CS acquisitions (Acquisition), using different AFs. Fully sampled (red), BART-CS (blue), and KLR-CS (green). The fully sampled reconstruction (AF=1), which does not differ between simulations and acquisitions, has been repeated to facilitate figure reading.

**Fig. 5:**
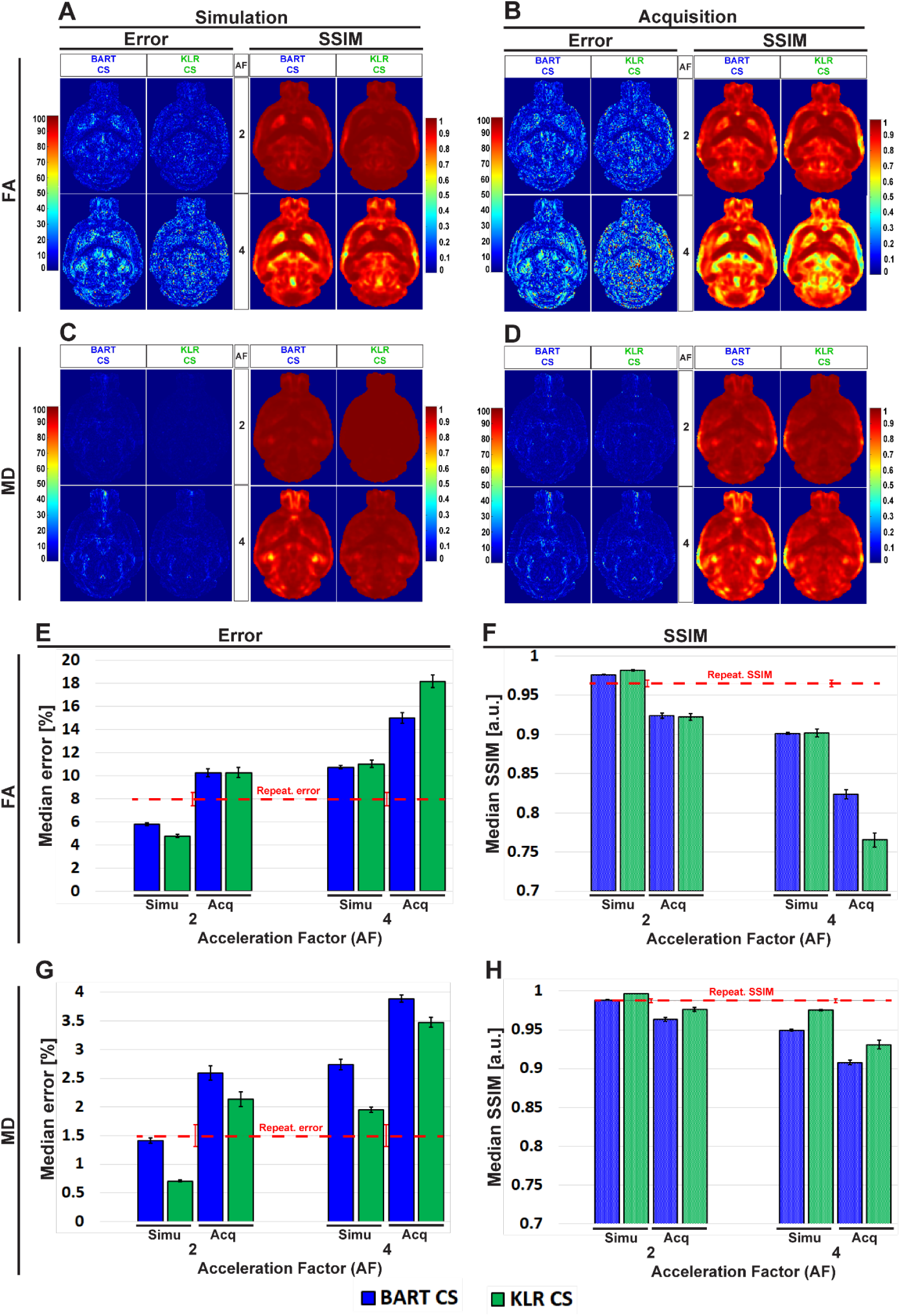
Error and SSIM, derived from FA and MD maps obtained using BART-CS and KLR-CS, for simulated undersamplings and for CS acquisitions. Error and SSIM for (**A,B**) FA and (**C,D**) MD maps and corresponding full 3D brain, (**E,G**) median error and (**F,H**) median SSIM obtained using BART-CS (blue) and KLR-CS (green) reconstructions, for the full 3D brain, and using fully sampled data as the reference. The left column **(A,C,E,G)**corresponds to retrospective undersampling (Simulation), and the right one **(B,D,F,H)**to CS acquisitions (Acquisition). The repeatability error and SSIM calculated from three repetitions of a fully sampled acquisition are represented as red dashed lines. Results expressed as mean ± standard deviation across animals (n=6).

We surprisingly observed that the error increased and SSIM decreased in acquisitions compared with simulations, for all AF values and for both FA and MD. For AF=2 and KLR-CS, the FA error increased from 4.79±0.12% in simulations to 10.28±0.44% in acquisitions (*p*<0.0001), and the FA SSIM decreased from 0.981±0.001% to 0.922±0.004% (*p*<0.0001). Similarly, the MD error increased from 0.71±0.02% in simulations to 2.13±0.12% in acquisitions (*p*<0.0001), and the MD SSIM decreased from 0.997±0.001% to 0.977±0.002% (*p*<0.0001). As the errors in acquisitions are surprisingly much larger than that observed in simulations, we explored the different contributions to these errors to ensure that the proposed acquisition method was working as intended.

First, the repetition noise was evaluated. The FS and the AF=2 DWI datasets were acquired three times on the same fixed brain. The comparison of the three FS datasets between themselves had a median error of 7.97±0.57% and a median SSIM of 0.965±0.004 for FA and 1.50±0.19% and 0.987±0.003 for MD (Supp. Fig. 5). These repetition errors are shown in Fig. 5 as a red dashed line, and they contribute to about two thirds of the observed median error.

Second, we evaluated the difference in the resonance frequency drift during a FS acquisition (13h37min) and during an AF=2 acquisition (6h49min). We observed drifts of 33.9±13.6 and 10.9±5.1 Hz, respectively. A part of this frequency drift induces a spatial shift, which is corrected during post-processing by image registration (0.6 pixels and 0.3 pixels for our two experimental conditions). The rest of the frequency drift may not be corrected for and alters the image. Because of the acquisition durations, we expected the FS data to be more affected than the CS data. To evaluate this contribution, we simulated a spatial drift and computed the median error and the SSIM between the FA and MD maps obtained with and without simulated drift. We observed an error that represents about 10% of the error between the FS and CS acquisitions (Fig. 5).

Third, we evaluated the change in peak signal-to-noise ratio (pSNR) between the FS and the CS acquisitions. For AF=2, the average pSNRs across diffusion directions of FS, CS simulations, and CS acquisitions were 318.0±26.9, 179.3±15.1, and 155.9±14.1, respectively. The 13% reduction in pSNR between CS simulations and CS acquisitions (*p*<0.0001) contributes to increasing the error on acquired FA and MD maps. In addition, we observed strong correlations between pSNR and the median error as a function of the diffusion direction: −0.7144 (p<0.0001) and −0.6785 (p<0.0001) for BART-CS and KLR-CS acquisitions, respectively. The noise level was also independent of the direction, suggesting that it remains stable when the undersampling pattern changes, as expected.

To evaluate the utility of the proposed CS approach, we considered two additional metrics. First, as all the reported metrics so far were obtained at the pixel level for the whole brain, we considered instead the error and the SSIM within two ROIs: the corpus callosum, a white matter ROI, and the caudate putamen, a gray matter ROI. This approach corresponds to what is usually performed to report diffusion-based estimates. Regarding the change in absolute value, the ROI-level error was lower than the pixel-level error reported in Fig. 5 for MD of gray matter and FA of white matter regions, and it was higher for MD of white matter and in some cases for FA of gray matter regions. In addition, we observed that KLR-CS almost always outperforms BART-CS in simulations and acquisitions (see Supp. Fig. 6 for details).

Second, we considered white-matter tracts. The main shape of the anterior commissure is preserved in both simulations and acquisitions (Fig. 6A-D). Both BART-CS and KLR-CS methods did not show significant differences in the mean fiber length in simulations and acquisitions to their corresponding WT and *MAP6*^+/-^ references for AF=2 (*p*>0.05 for all) (Fig. 6E,F). However, for AF=4, CS acquisitions showed a reduction in mean fiber length for both WT (−11.9%, unpaired *t*-test, *p*=0.0059) and *MAP6*^+/-^ (−5.9%, unpaired *t*-test *p*=0.0424) mice when compared with CS simulations. With our small dataset, the difference in fiber length between WT and *MAP6*^+/-^ mice, visible for the FS and AF=2 data, is no longer significant for AF=4. A similar analysis was made for the fornix, a tract with increased complexity, given its wide distribution in the 3D space and higher levels of defasciculation than the anterior commissure. As previously found, the main shape of the fornix and the mean fiber length were conserved by both CS methods (see Supp. Fig. 7 for details). Moreover, KLR-CS showed a smaller variance on the mean fiber lengths than BART-CS (*p*<0.05, considering data from the anterior commissure and fornix). These results suggest that AF=2 seems to have no impact on fiber tract analysis, and AF=4 has a small but acceptable impact. In addition, KLR-CS appears as the most robust method.

**Fig. 6:**
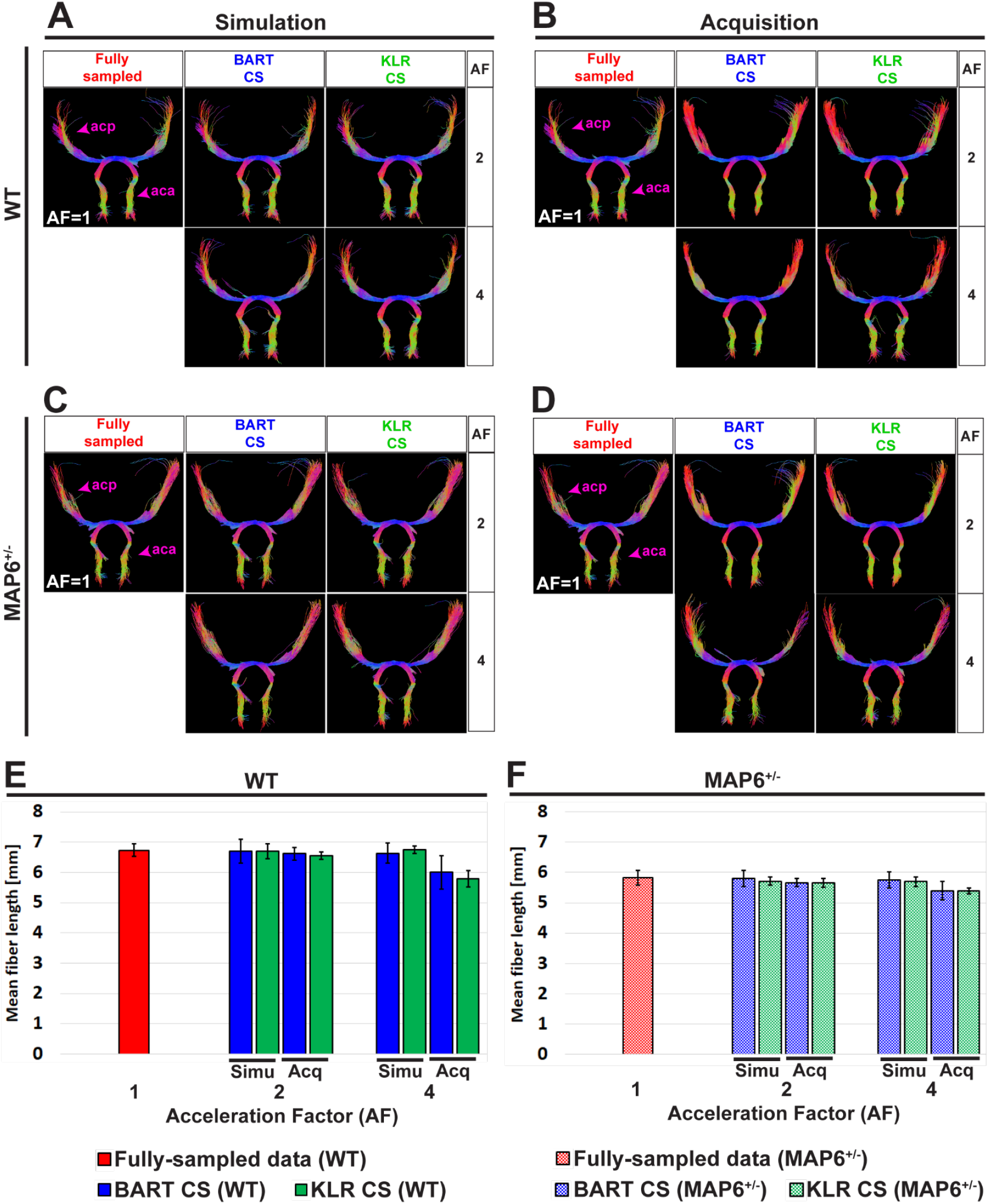
Tractography of the anterior commissure (ac) from fully sampled data, after simulated undersampling, and for CS acquisitions, using BART-CS and KLR-CS reconstructions. Anterior commissure (ac) of a (**A,B**) WT and a (**C,D**) *MAP6*^+/-^ mouse from fully sampled (red), BART-CS (blue), and KLR-CS (green) reconstructions, using different AFs. Mean fiber length of the ac of (**E**) WT and (**F**) *MAP6*^+/-^ mice. The left column **(A,C,E)** shows results obtained with simulations and the right column **(B,D,F)** with acquisitions. The anterior commissure from fully sampled data is repeated between simulation and acquisition columns, to facilitate figure reading. Results are expressed as mean ± standard deviation across animals (n=3 per group). acp: ac posterior part; aca: ac anterior part.

## 4. Discussion

In this study, we implemented an original 3D Spin-Echo CS acquisition method for preclinical DWI imaging, with the possibility of using the same or different sampling masks per diffusion direction. Two reconstruction algorithms were evaluated: BART-CS and the more recently proposed KLR-CS approach. After an optimization step based on retrospective undersampling, FA and MD maps, as well as tractographies, were produced based on CS data acquired at 9.4T using fixed mouse brains. The two reconstruction protocols yielded robust results for a reduction in scan time up to 4, with a clear advantage for KLR-CS. When FA and MD were estimated in a white-matter ROI, the difference between FS and KLR-CS reconstructions remained below 3.9% and 7.0%, for AF=2 and 4, respectively. Regarding tracts, a reduction in scan time up to 4 produced only minor faults, even in a complex structure such as the fornix.

Moreover, we used a definition of AF that differs from the one used in previous studies^21^: AF was considered here as the effective reduction in scan time. Therefore, as we chose to fully sample the b0 images, the AF used per diffusion direction (AF_diff_) was higher. For instance, to reach an effective AF of 4, we used an AF_diff_ of about 5.71. To facilitate the comparison with previous CS DWI studies, Supp. Table 1 provides a summary of simulation and acquisition parameters in this and previous studies, including both AF and AF_diff_.

As the published KLR-CS requires a fully sampled dataset for regularization purposes^21,22^, we proposed two approaches to deal with acquired complex data and chose, based on simulations, the KLR-CS based on LRP maps, which, with AF ≤6, outperformed BART-CS. Further improvements may be proposed. First, coil sensitivity maps calculated by robust algorithms such as ESPIRiT^30^, important for the quality of BART-CS reconstructions, are not used in the current implementation of KLR-CS, which uses simpler sensitivity maps. Second, the size of the central part of the k-space was set based on Zhang et al’s study and therefore on their magnitude-filtered approach^21^. This size, important for the learning step of the KLR-CS, could be optimized for the proposed LRP approach. To further improve the quality of the reconstruction, the LRP KLR-CS approach could also be challenged by different theoretical frameworks, such as manifold-modeled signal recovery algorithms^41^ or model-based methods with spatial and parametric constraints^42^. Further work is required to evaluate the combination of CS with parallel imaging^9^. Note that additional regularization along the diffusion direction could be performed with the BART toolbox^29^. This would require further tuning of the CS parameters but could improve the BART results.

With our data and choice of parameters, both BART-CS and KLR-CS reconstructions yielded lower errors than that reported by Zhang et al.’s study^21^, which was obtained on simulated CS data, using equivalent ROIs but with magnitude-filtered KLR-CS (Supp. Fig. 6 in our study; Fig. 3 in Zhang et al.’s article^21^). The difference could be related to the difference in acquisition conditions (Supp. Table 1), but lower errors remain nevertheless encouraging. Regarding major white-matter tracts, the main shape is always conserved in both simple (anterior commissure) and more complex (fornix) tracts when using CS acquisitions, for all AFs studied. The appearance of some false-positive fibers, especially in the fornix, can partially be controlled by the addition of exclusion ROIs. For AF=4 and both CS approaches, further adjustment of tractography parameters, use of deep-learning-based versions of CS^43–46^, and possibly introducing priors^47,48^ could help improve the reconstruction, but they have not yet been applied to preclinical data. To allow comparisons with other reconstruction methods, the data acquired in this study have been made available.

The switch from simulations to acquisitions increased the observed error and, consequently, the observed SSIM was decreased. Additional fluctuations with diffusion directions were also observed. We, therefore, analyzed the different contributions to this increase in error and fluctuations.

First, we characterized the repeatability (Supp. Fig. 5). Indeed, when using simulations, the values in the undersampled dataset are exactly the same as in the FS datasets. In the case of acquisitions, the values in the undersampled dataset differ from that in the FS dataset. We observed that this repetition error (7.97±0.57% for FA, 1.50±0.19% for MD) was about twice the error of the KLR-CS reconstruction in simulations (AF=2; 4.79±0.12% for FA, 0.71±0.02% for MD, Fig. 5). The repetition noise appears as the main contribution to the difference between acquisitions and simulations.

Second, there are differences in acquisition conditions between simulated and acquired data. These differences include the effect of the frequency drift (about 10% of the error) and the small reduction in pSNR (about 13%), which also contributes to increasing the error on the derived parameters (FA, MD, and tracts). To limit the impact of the frequency drift, a navigator could be introduced in the MRI sequence^17^. Note that, concerning the frequency drift, the FS dataset is more blurred than the undersampled ones. The “error” is therefore not always in the CS datasets.

Third, the spatial correction, applied to limit the effect of the frequency drift during acquisition, could be improved. Here, we used an algorithm to modify the phase in k-space and thereby shift the image, to linearly register DWI images to the reference. However, the effect of the difference in acquisition times between FS and CS acquisitions on image distortion remains to be evaluated and, if necessary, corrected for. In addition, the MRtrix3 probabilistic reconstruction can add some noise to the fiber tract reconstructions. Fifty repetitions of the MRtrix3 reconstruction on the same dataset led to 1% repetition variability on mean fiber length — a small but not negligible contribution to the error.

To summarize, estimates made on acquired data are in good agreement with simulations, in which only CS reconstruction noise is present. Altogether, our results suggest that a reduction in scan time by a factor of 4 is acceptable in our experimental setting, in line with previous preclinical^16,21^ and clinical results^42,49^. Further analysis includes increasing the number of directions and the spatial resolution^6,16^ to achieve higher AFs without degrading image quality^50^ and limiting the increase in scan time as the blurring for KLR-CS is in both the spatial and diffusion directions. Indeed, this reconstruction approach should benefit from a larger learning set in both spatial and diffusion dimensions. A concurrent optimization of the spatial resolution and the number of diffusion directions could thus be performed^16^.

There were some limitations to the study. We used only three animals per group, given that our objective was not biological but methodological; this small number limits the power of the statistical analysis. Further studies are required to fully characterize this mouse model. Similarly, we focused on a few ROI and a few tracts, in parallel to 3D FA and MD maps. More tracts could be analyzed based on the data that have been shared (see ‘Data and code availability’). Moreover, acquisitions were performed using only one MRI scanner. A multicentric study would be more appropriate to estimate acquisition and reconstruction errors as the choice of hardware may affect the outcomes^51^.

In conclusion, LPR KLR-CS reconstruction of undersampled preclinical DWI data seems a robust approach to reducing scan time while maintaining accurate estimates of FA, MD, and tracts.

## Abbreviations

AF: Acceleration Factor
b0: b~0 s/mm^2^ images (baseline images acquired without diffusion weighting)
CS: Compressed Sensing
DWI: Diffusion-Weighted Imaging
FA: Fractional Anisotropy
FS: Fully sampled
KLR: Kernel Low-Rank
LRP: Low-resolution phase
MAP6: Microtubule-associated protein 6
MD: Mean Diffusivity
pSNR: Peak signal-to-noise ratio
ROI: Region of interest
SNR: Signal-to-noise ratio
SSIM: Structural Similarity Index Measure
WT: Wild type

## Acknowledgements

We are particularly grateful to Dr. Leslie Ying, Ukash Nakarmi and Chaoyi Zhang at the University of New York at Buffalo, for their toolbox for KLR Compressed Sensing reconstruction and the support to handle it and to Sylvie Gory-Fauré for providing the WT and *MAP6*^+/-^ mice.

## Data and code availability

Raw data and code are available at https://github.com/nifm-gin/compressedSensing (content will be made available after publication).

## Supplementary Materials

Supp. Tables 1 to 2

Supp. Figs. 1 to 7

