## Supplementary Tables and Figures for "Evaluation of Kernel Low-Rank Compressed Sensing in preclinical Diffusion Magnetic Resonance Imaging"

### Supplementary material

Diego ALVES RODRIGUES DE SOUZA[1], Hervé MATHIEU[1, 2], Jean-Christophe DELOULME[1#], Emmanuel L. BARBIER[1, 2#*]

[1]Univ. Grenoble Alpes, Inserm, U1216, Grenoble Institut Neurosciences, Grenoble, France.

[2]Univ. Grenoble Alpes, Inserm, US17, CNRS, UAR 3552, CHU Grenoble Alpes, IRMaGe, Grenoble, France.

#Co-Last authors

*Corresponding author

Emmanuel L. BARBIER
Grenoble Institut des Neurosciences – U1216
Team «Functional Neuroimaging and Brain Perfusion»
Chemin Fortuné Ferrini
38700 La Tronche


| Image type | Magnetic field (T) / gradient strengh (mT/m) / receive coil type | Resolution (µm$^3$) / matrix size | TR / TE (ms) | b-value (s/mm$^2$) | b0 images | Diffusion directions | Study type | $AF_{diff}$ | AF | Scan time (h) |
|---|---|---|---|---|---|---|---|---|---|---|
| DWI (3D Spin Echo) [this study] | 9.4 / 660 / 4-channel cryocoil | 100 / 180x90x110 | 150 / 19 | 3000 | 3 | 30 | Reference | 1.00 | 1.0 | 13.6 |
| | | | | | | | CS simulation | 2.22 | 2.0 | 6.8 |
| | | | | | | | | 3.75 | 3.0 | 4.5 |
| | | | | | | | | 5.71 | 4.0 | 3.4 |
| | | | | | | | | 12.00 | 6.0 | 2.3 |
| | | | | | | | CS acquisition | 2.22 | 2.0 | 6.8 |
| | | | | | | | | 5.71 | 4.0 | 3.4 |
| DWI (3D DW-GRASE) [Zhang et al's study[21]] | 7 / - / 4-channel cryocoil | 100 / 128x104x116 | 500 / 33 | 2000 | 2 | 30 | Reference | 1.0 | 1.00 | 8.0 |
| | | | | | | | CS simulation | 2.0* | 1.88* | 4.3* |
| | | | | | | | | 4.0* | 3.37* | 2.4* |
| | | | | | | | | 6.0* | 4.57* | 1.8* |
| | | | | | | | | 8.0* | 5.56* | 1.4* |
| DWI (3D Spin Echo) [Wang et al's study[16]] | 9.4 / 2000 / - | 45 / 420x256x256 | 100 / 12.7 | 4000 | 5 | 46 | Reference | 1.0 | 1.0 | 92.8 |
| | | | | | | | CS acquisition | 4.0 | 4.0 | 23.2 |
| | | | | | | | | 5.12 | 5.12 | 18.2 |
| | | | | | | | | 6.4 | 6.4 | 14.5 |
| | | | | | | | | 8.0 | 8.0 | 11.6 |

**Supp. Table 1: Main scan parameters for this study and two reference studies**

*: parameter value not explicitly mentioned in the article, but whose value could be derived

| Neuronal Tract | ROI type | Orientation | Coordinate [mm] (B: Bregma, L: Lateral) | Target region | Figure number (Atlas Franklin & Paxinos) |
|---|---|---|---|---|---|
| Anterior commissure (ac) | AND | Sagittal | 0.24 L | ac | 103 |
| | AND | Sagittal | -0.24 L | ac | - |
| Fornix (f) | AND | Horizontal | -3.60 B | f | 144 |
| | AND | Coronal | -0.58 B | f | 36 |
| | NOT | Coronal | -0.82 B | sm | 38 |

**Supp. Table 2: ROIs used to obtain the anterior commissure and the fornix tracts.**

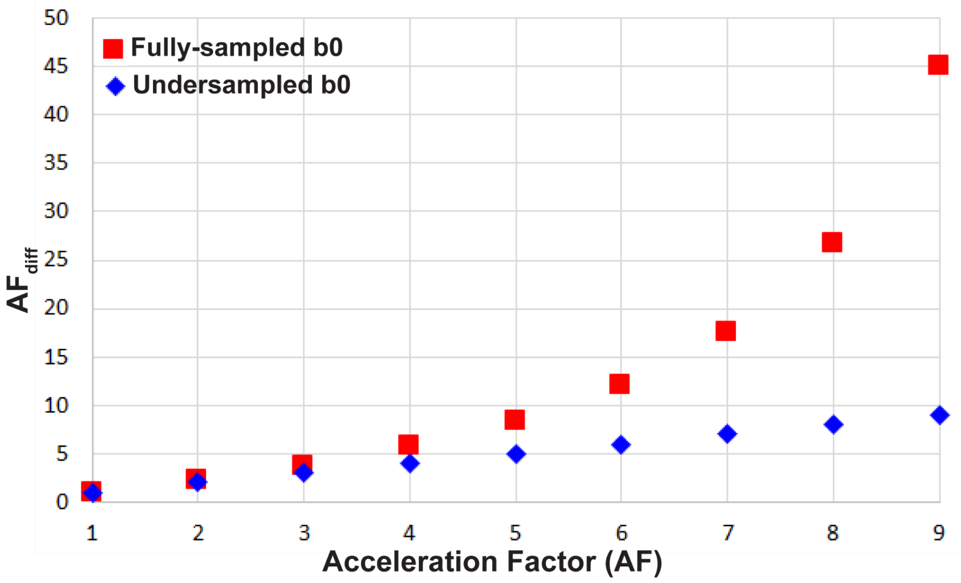


**Supp. Fig. 1: AF for diffusion directions ($AF_{diff}$) as a function of AF (the global acceleration factor).**

$AF_{diff}$ as a function of AF for a DWI acquisition with 3 b0 and 30 diffusion direction images, considering fully sampled b0 (red dots) and undersampled b0 (blue dots) images. The details of these parameters and their relations are as follows:

$$AF = \frac{(n_{b0} + n_{diff}) \cdot (AF_{b0} \cdot AF_{diff})}{(n_{b0} \cdot AF_{diff} + n_{diff} \cdot AF_{b0})},$$

where AF, $AF_{diff}$, $AF_{b0}$, $n_{diff}$ and $n_{b0}$ are the chosen AF of the entire acquisition, the AF for the diffusion directions, the AF for the b0 images, the number of diffusion directions and the number of b0 images, respectively. In the case of fully sampled b0 images (red dots), AF is obtained by undersampling the diffusion directions only. Thus, when AF > 1, $AF_{diff}$ is higher than the chosen AF. Moreover, when b0 images are fully sampled, $AF_{diff}$ may be obtained by:

$$AF_{diff} = n_{diff} \cdot AF / [(n_{b0} + n_{diff}) - (n_{b0} \cdot AF)].$$

When b0 images are undersampled (blue dots), the acceleration is uniformly distributed over all DWI volumes, and $AF_{diff}$ = AF.

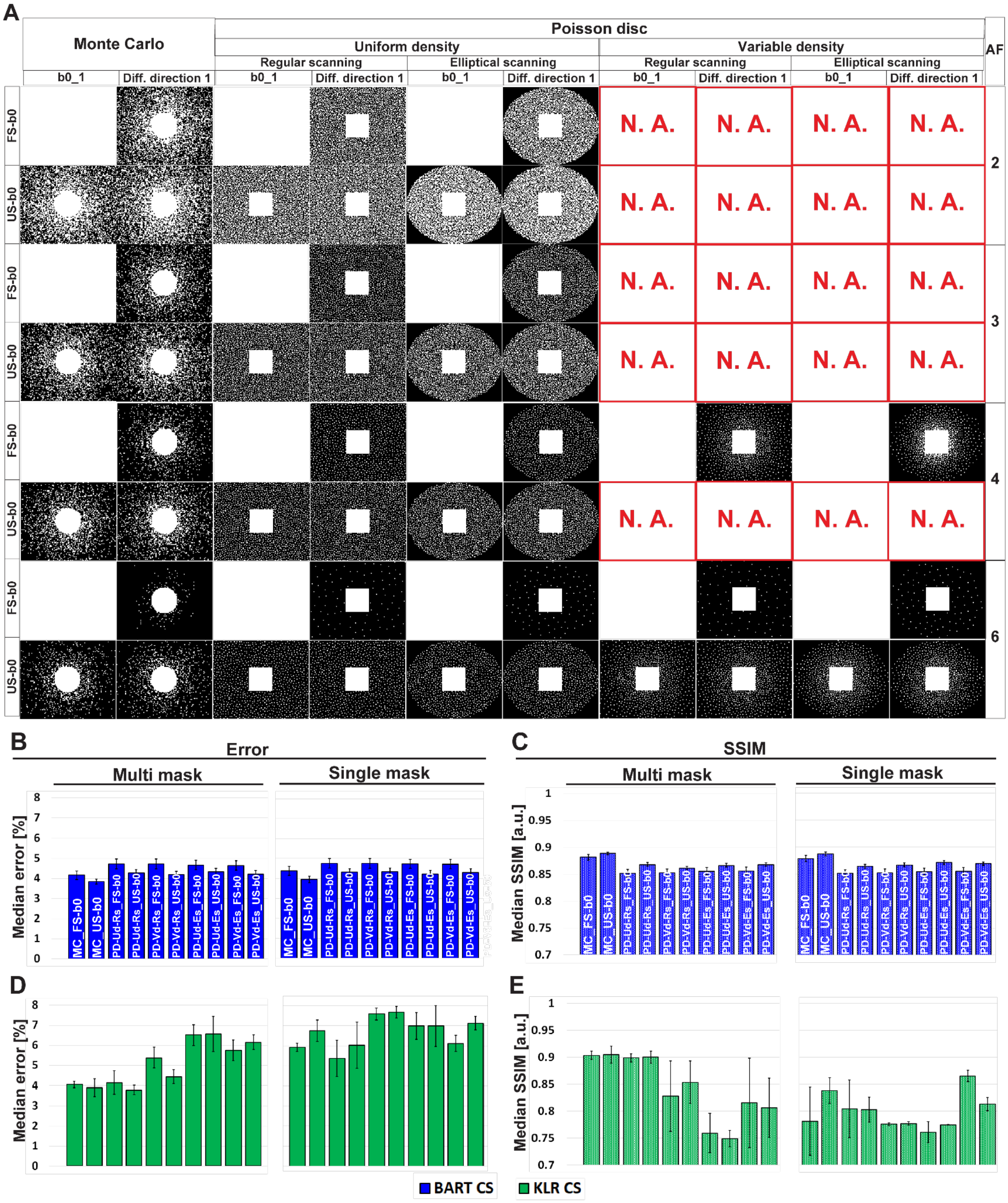


**Supp. Fig. 2: Optimization of the undersampling strategy**

(**A**) Examples of undersampling patterns for AF=2 to 6. Corresponding (**B,D**) error and (**C,E**) SSIM values, using MD maps and AF=6. **(B,C)** correspond to BART-CS and **(D,E)** to KLR-CS reconstruction. Results are expressed as mean ± standard deviation across animals (n=3)

Columns show different Poisson-disc and Monte Carlo undersampling patterns and their resulting masks for the first b0 image and the first diffusion direction. Rows correspond to AF (between 2 and 6) and sampling mode of b0 images (FS: fully sampled; US: undersampled). Note that only phase encoding directions are undersampled. "N.A." stands for "not available" (a mask cannot be generated in these specific conditions). A total of 20 possible combinations of undersampling patterns per AF was evaluated (Poisson disc: fully sampled or undersampled b0 / Uniform or variable density / single or multi-mask mode / common or elliptical scanning; Monte Carlo: fully sampled or undersampled b0 / single or multi-mask mode). The naming of the bar graph: MC: Monte Carlo, PD: Poisson disc, Ud: Uniform density, Vd: variable density, Rs: regular scanning, Es: elliptical scanning.

With the KLR-CS reconstruction, the multi-mask undersampling outperformed the single-mask undersampling (D,E). For instance, for a Monte-Carlo with FS b0s undersampling scheme ("MC_FS-b0" bars) and AF=6, the error reduced from 5.92±0.21 to 4.06±0.16% (*p*=0.0054), and the SSIM increased from 0.781±0.063 to 0.917±0.020 (*p*=0.0479) when replacing the single-mask undersampling with a multi-mask. With the BART-CS reconstruction, single and multi-mask undersampling has no effect on error or SSIM (B,C).

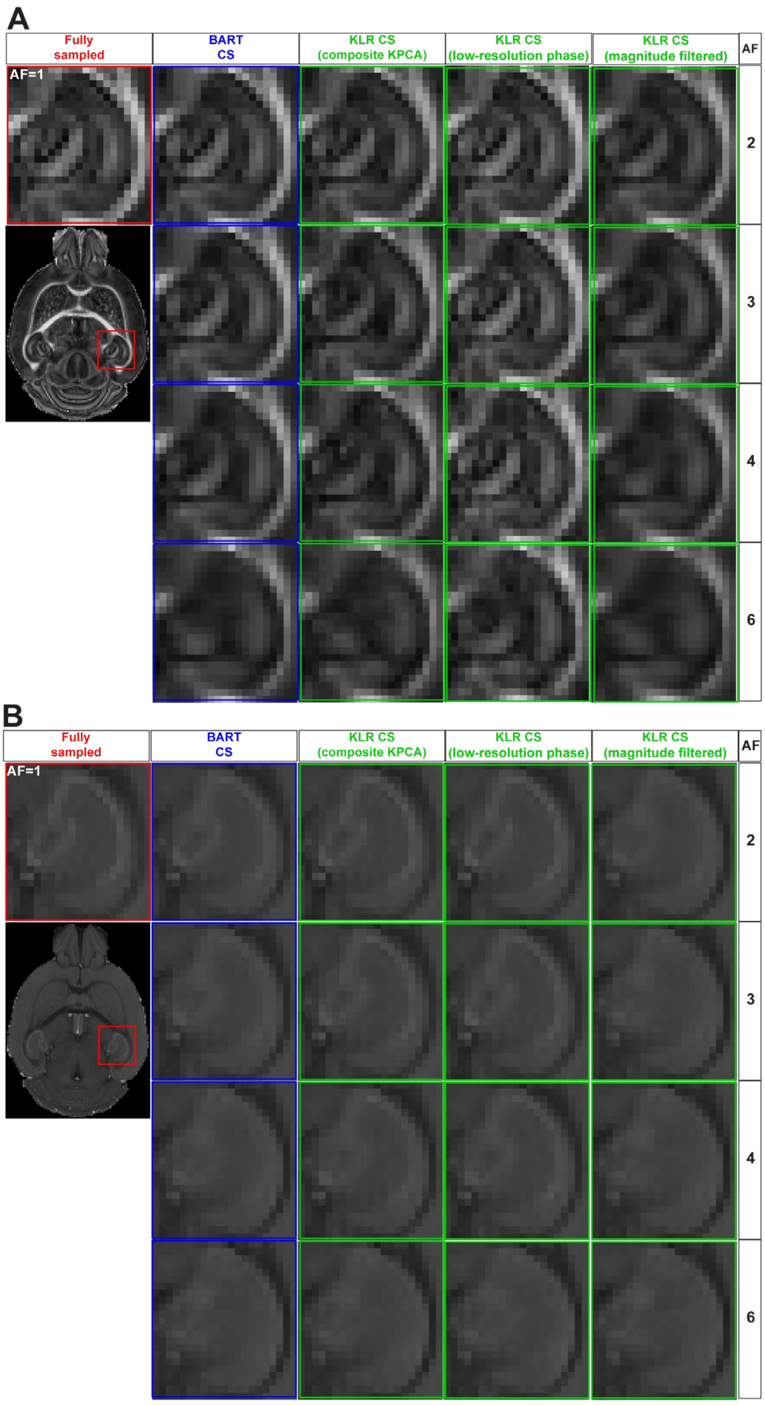


**Supp. Fig. 3: FA and MD maps from fully sampled data and after simulated undersampling, reconstructed using BART-CS, KLR-CS based on composite KPCA, KLR-CS with low-resolution-phase (LRP) maps, and magnitude-filtered KLR-CS.**

(**A**) FA and (**B**) MD maps from a zoom on the hippocampus, obtained from simulated reconstructions by BART (blue) and three different implementations of KLR-CS (green), for different AFs. A reconstruction from a fully sampled dataset (red) is shown as the reference.

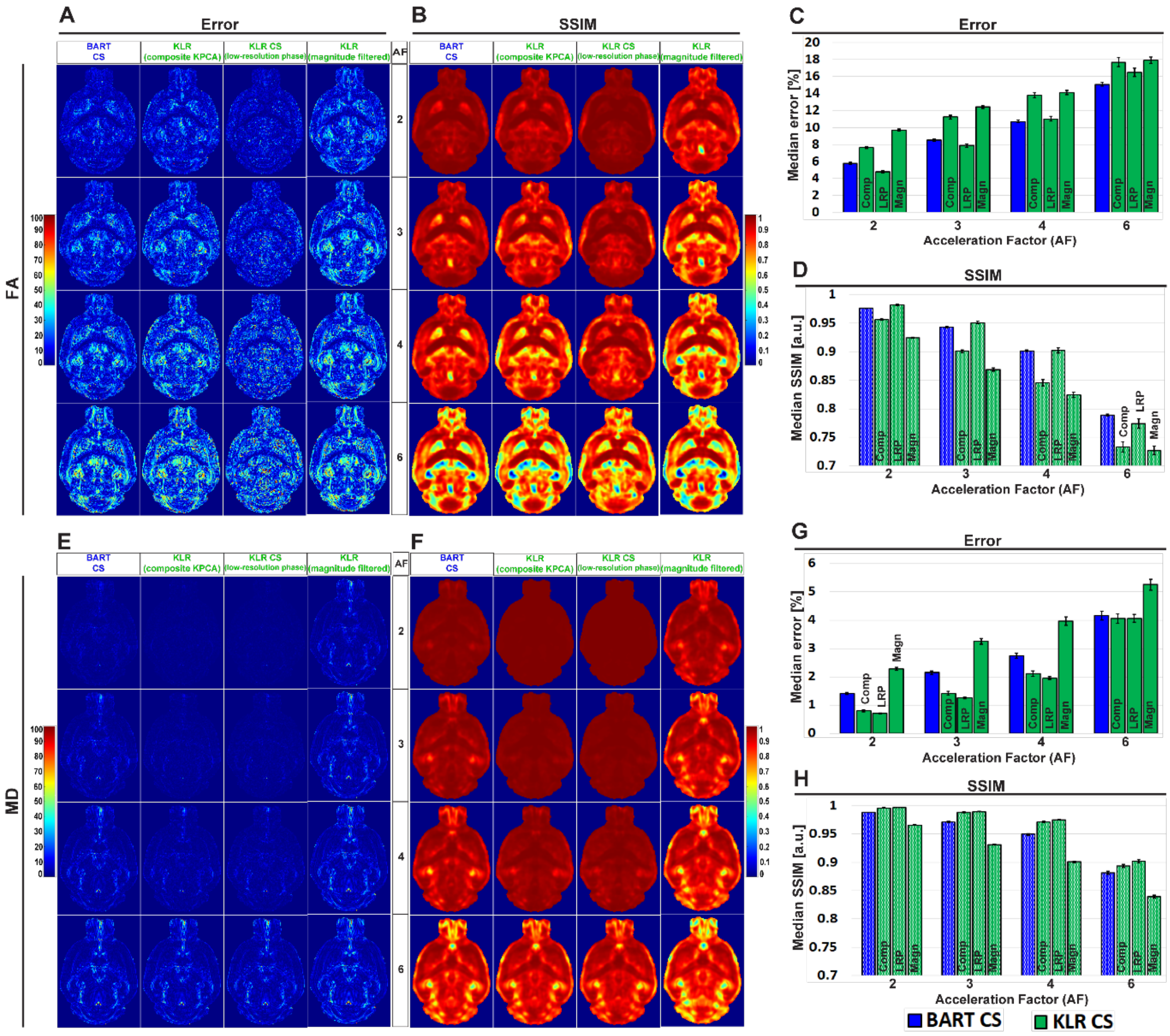


**Supp. Fig. 4: Error and SSIM on FA and MD, from simulated undersampling reconstructed using BART-CS, KLR-CS based on composite KPCA, KLR-CS with low-resolution-phase (LRP) maps, and magnitude-filtered KLR-CS.**

Error and SSIM of (**A,B**) FA and (**E,F**) MD maps using BART-CS and three different implementations of KLR-CS, for different AFs. Full 3D brain values of (**C,G**) median error and (**D,H**) median SSIM, using the fully sampled data as the reference. Results expressed as mean ± standard deviation across the animals (n=3). The three KLR-CS implementations are KLR-CS based on composite KPCA (Comp), KLR-CS using low-resolution-phase (LRP) maps, and magnitude-filtered KLR-CS (Magn).

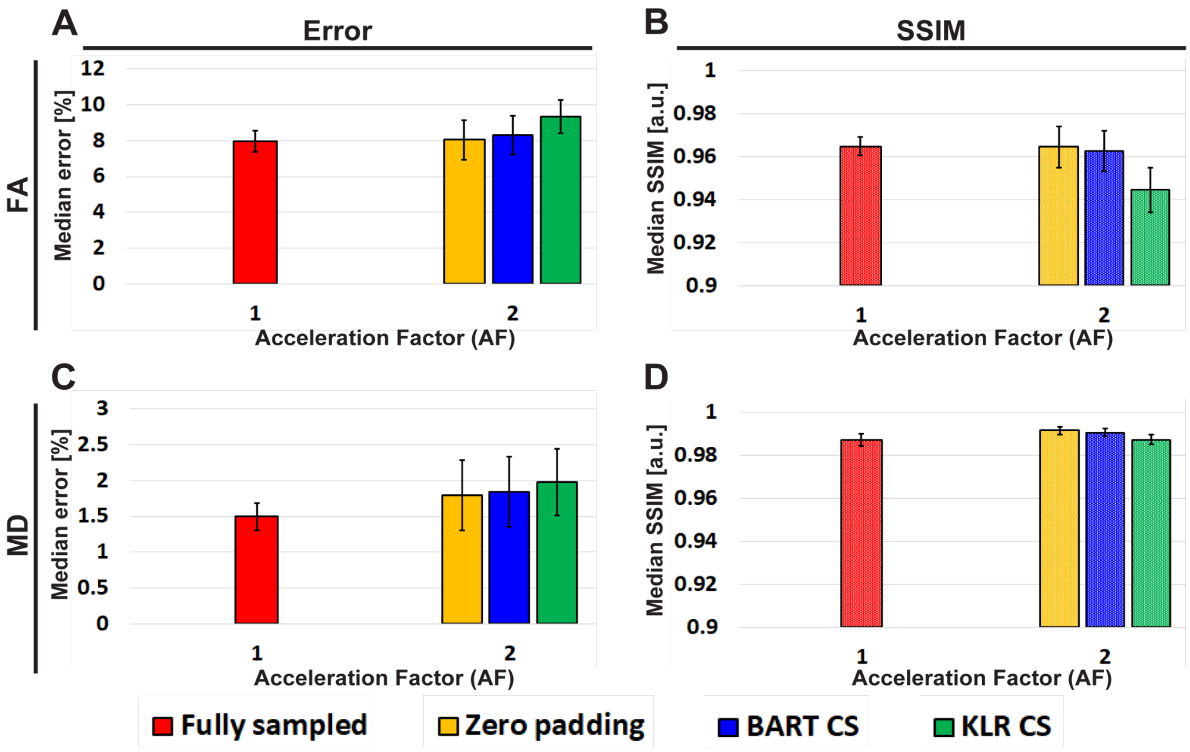


**Supp. Fig. 5: Repeatability error and SSIM on FA and MD, derived from fully sampled and CS acquisitions.**

Repeatability error and SSIM of (**A,B**) FA and (**C,D**) MD maps for fully sampled and reconstructions by zero padding, BART-CS and KLR-CS of undersampled acquisitions. The repeatability metrics were calculated from three repetitions of a fully sampled and three repetitions of an AF=2 undersampled acquisition and the same sample. The repeatability metrics of fully sampled data were used as the references of Fig. 5. The repeatability noise on fully sampled and undersampled data were comparable, except for FA maps of the KLR-CS method, which exhibit a slightly decreased SSIM. Results are expressed as mean ± standard deviation across repetitions (n=3).

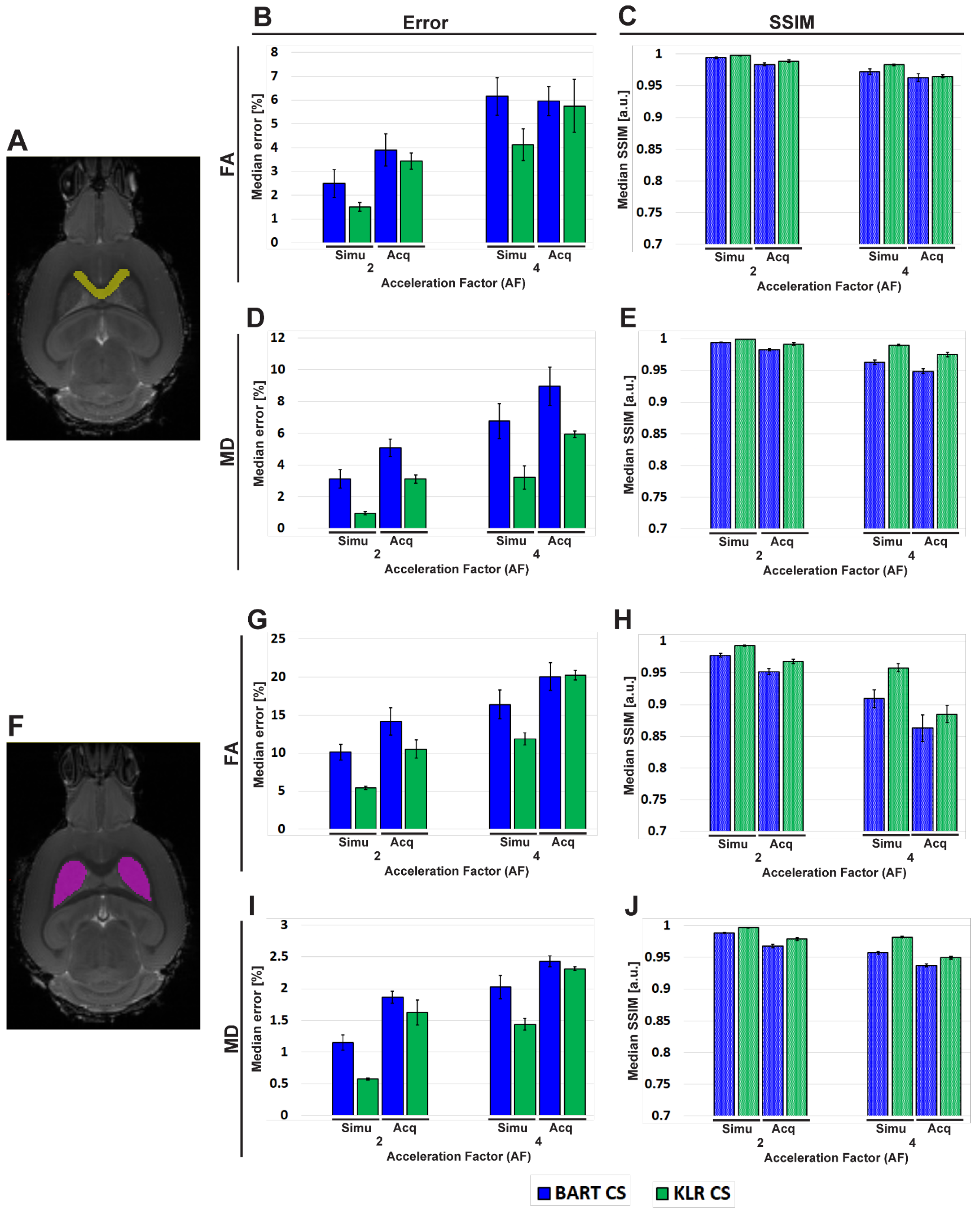


**Supp. Fig. 6: FA and MD measured in two regions of interest**

Analysis of median error and SSIM within specific ROIs: (**A**) corpus callosum (cc) and (**F**) caudate putamen (CPu). The first column **(B,D,G,I)** correspond to the error and the second **(C,E,H,J)** to the SSIM of (**B,C**) FA and (**D,E**) MD maps of cc, as well as for (**G,H**) FA and (**I,J**)

MD maps of CPu. Both metrics were obtained using reconstructions by BART-CS (blue) and KLR-CS (green) from CS acquisitions and using fully sampled data as the reference. Results are expressed as mean ± standard deviation across animals (n=3).

For FA maps of acquisitions with AF=2, the errors for the whole brain were respectively 10.27±0.35% and 10.28±0.44% for BART and KLR-CS (Fig. 5E), although for the cc (white matter) they were lower: 3.90±0.68% and 3.43±0.35%, respectively ($p$=0.0078 and $p$=0.0017) (B); and for the CPu (mainly gray matter) the errors were significantly higher only for BART-CS: 13.81±1.31% (BART-CS) and 10.55±1.15% (KLR-CS) ($p$=0.0477 and $p$=0.7274, respectively) (G). Yet, the inverse situation took place when analyzing MD maps of acquisitions under AF=2: the errors for the whole brain were respectively 2.59±0.12% and 2.13±0.12% for BART and KLR-CS (Fig. 5G), while for the cc (white matter) they were higher: 5.08±0.56% and 3.11±0.26%, respectively ($p$=0.0290 and $p$=0.0098) (D); and for the CPu (mainly gray matter) the errors were lower: 1.86±0.10% and 1.62±0.20% (I) for BART and KLR-CS, respectively ($p$=0.0107 and $p$=0.0166).

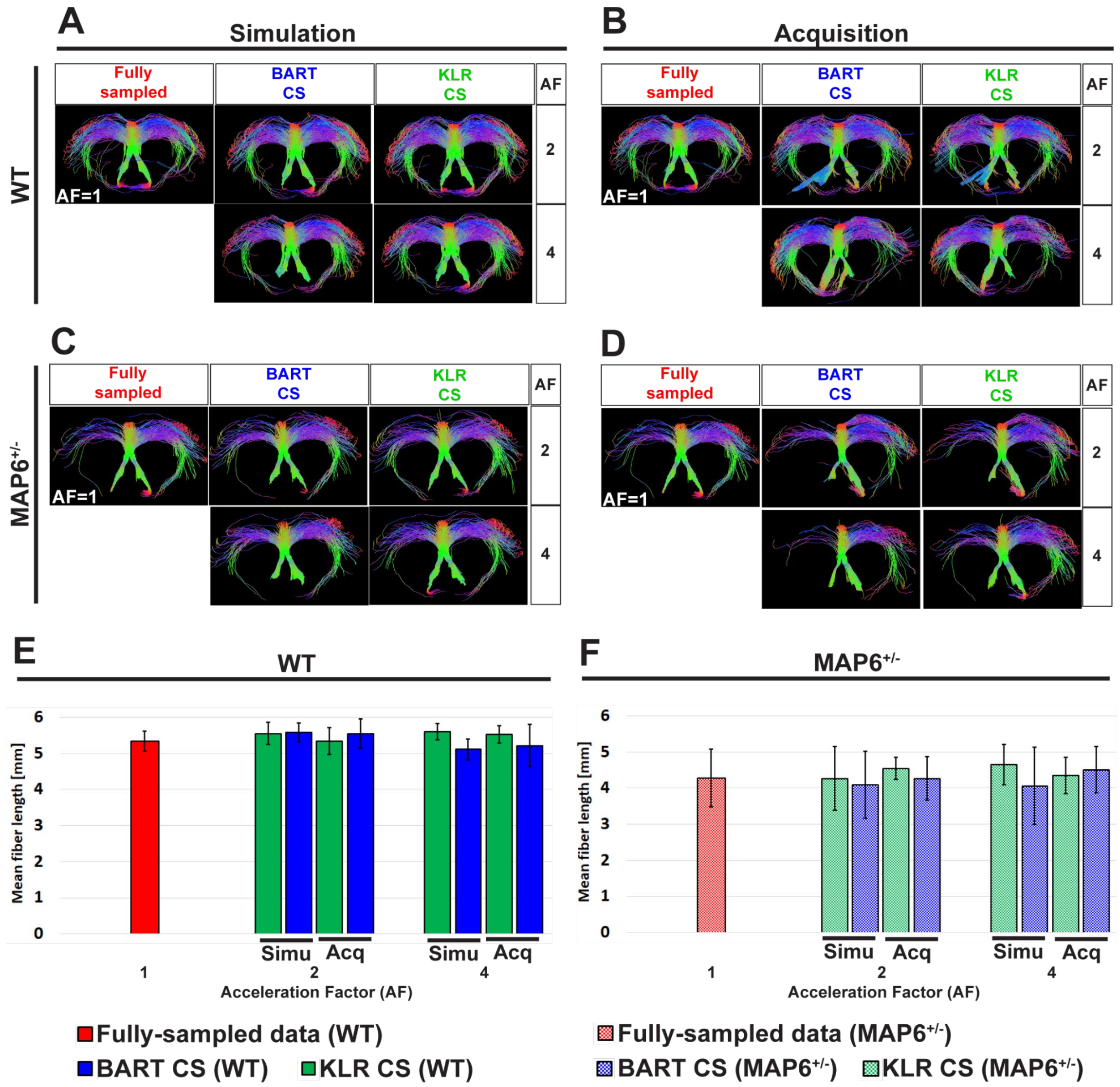


**Supp. Fig. 7: Tractography of the fornix from fully sampled data, simulated undersampling, and CS acquisitions.**

Fornix of a (**A,B**) WT and a (**C,D**) *MAP6*$^{+/-}$ mouse from a fully sampled (red), BART-CS (blue), and KLR-CS (green) reconstructions, using different AFs. Mean fiber length of the fornix of (**E**) WT and (**F**) *MAP6*$^{+/-}$ mice. The left column **(A,C,E)** stands for results obtained with simulated undersampling and the right one **(B,D,F)** with CS acquisitions. The fornix from fully sampled data is repeated between simulation and acquisition columns, to facilitate figure reading. Results are expressed as mean ± standard deviation across animals (n=3 per group).

The fornix is a tract with increased complexity, given its wide distribution in the 3D space and higher levels of defasciculation than the anterior commissure. As for the ac, the main shape of the fornix was conserved by both CS methods in real acquisitions and AF=2 w.r.t. the fully sampled data. For AF=2, one can see the presence of false positive fibers at the level of the post-commissural ventral fornix terminus, if the same ROIs for filtering this tract are used. These false positives were easily removed by a supplementary ROI (not shown). For AF=4, the presence of false negatives may readily be seen in (A,B), at the level of both post-commissural dorsal part and left part of the post-commissural ventral fornix terminus. This yielded misleading differences between WT and *MAP6*$^{+/-}$. The mean fiber length of CS simulations and acquisitions showed no significant difference when directly compared to their fully sampled references for WT and *MAP6*$^{+/-}$ ($p$>0.05 for all) (E,F). The mean fiber length of WT and *MAP6*$^{+/-}$ from fully sampled data did not show a significant difference for the fornix (unpaired $t$-test, $p$=0.1520). The same held when this comparison was made for CS simulations and acquisitions, in such a way that both BART and KLR-CS reproduced the fully sampled pattern, for AF=2 and 4 (unpaired $t$-test, $p$>0.05).
